# Multi-omics Reveals Divergent Endothelial Molecular Responses to New and Old World Hantaviruses

**DOI:** 10.64898/2026.09.11.751079

**Authors:** Yennifer Delgado, Anne K. Zaiss, Immy A. Ashley, Arjit Vijey Jeyachandran, Shipra Sharma, Vrithika Vaithi, Declan M. Winters, Andrea O. Chan, Pritha Sankar, Sophie F. Blanc, Charles O. Olwal, Justin Selby, Valerie Vandersall, Sara K. Makanani, Evan T. Wynn, Jocelyn J. Ni, Kareem Alba, Sandra Papazian, Brian Nguyen, Ashok Kumar, Shantanu Joshi, Gustavo Garcia, Nevan J. Krogan, Robert Damoiseaux, Vaithilingaraja Arumugaswami, Mehdi Bouhaddou

## Abstract

Hantaviruses cause vascular leakage syndromes that vary in clinical manifestation and severity. Although tissue tropism contributes to these differences, both Old and New World hantaviruses infect endothelial cells, where species-specific disease phenotypes remain poorly understood. Here we integrated time-resolved global RNA sequencing, mass spectrometry proteomics, and phosphoproteomics of human endothelial cells infected with New World Andes virus (ANDV) or Old World Hantaan virus (HTNV). Despite equivalent early viral RNA and protein levels, ANDV elicited a stronger innate immune protein response, preceding its restriction while HTNV replication continued. At later stages of infection, HTNV induced downregulation of cytoskeletal and junctional protein phosphorylation, accompanied by visual disruption of cellular actin architecture. Additionally, ANDV induced heightened activity of ERBB-family kinases, whose chemical inhibition by neratinib and afatinib reduced viral replication. Together, these data define species-specific responses in endothelial cells, identify druggable host targets, and reveal mechanisms with relevance to divergent vascular leakage symptomology.

## INTRODUCTION

Hantaviruses are relatively uncommon globally, yet infections caused by certain species can result in severe disease with exceptionally high case-fatality rates. In humans, pathogenic hantaviruses cause two major vascular leakage syndromes: hemorrhagic fever with renal syndrome (HFRS), classically associated with Old World hantaviruses such as Hantaan virus (HTNV), and hantavirus cardiopulmonary syndrome (HCPS), associated with New World hantaviruses such as Andes virus (ANDV) and Sin Nombre virus (SNV)^1,2^. ANDV infection has a reported case fatality rate of ∼30-40%, whereas HTNV is ∼5-15%^2,3^. Hantaviruses are traditionally known to be zoonotic viruses, maintained primarily in small mammal reservoirs and transmitted to humans most commonly through exposure to excreta from infected rodents^1,2^. However, ANDV is unique among hantaviruses, since human-to-human transmission has been documented^4,5^. Although ANDV and HTNV share substantial genomic similarity, the mechanisms underlying their distinct clinical manifestations remain poorly understood.

Hantaviruses have recently re-emerged as a prominent global health concern. In 2025, HCPS caused the death of Betsy Arakawa in New Mexico, the wife of renowned actor Gene Hackman, and three other deaths in California^6,7^. More recently, on May 2, 2026, the World Health Organization (WHO) was notified of a cluster of severe respiratory illnesses aboard the MV Hondius cruise ship, which was carrying 147 passengers and crew from 23 countries on a voyage originating in Argentina^6^. The outbreak resulted in 13 confirmed or probable cases, several laboratory-confirmed as the New World hantavirus, ANDV, and three deaths. Despite research efforts, no effective prophylactic or therapeutic direct-acting or host-directed antiviral agents have been approved for hantaviruses in the United States, limiting the capacity to prevent or respond to lethal outbreaks.

HTNV and ANDV cause distinct clinical syndromes^1,7,8^ that differ in severity^9–13^. ANDV causes an especially severe pulmonary vascular leak syndrome characterized by rapid onset edema, hypoxemia, respiratory failure, and shock, contributing to its high case-fatality rate. Although both Old and New World hantaviruses have been shown to efficiently infect endothelial cells^12,14–16^, differences in cellular or tissue tropism are thought to contribute to divergent disease manifestations. For instance, HTNV can infect renal endothelial, epithelial, and glomerular cell populations^1,2^, whereas ANDV shows tropism for pulmonary and lymphatic microvascular endothelium^1,17^. However, their overlapping capacity to infect endothelial cells across multiple organs raises the possibility that virus-specific endothelial responses also shape their divergent disease phenotypes.

Hantaviruses are enveloped, negative-sense RNA viruses in the family *Hantaviridae*, possessing a tripartite genome comprising the large (L), medium (M) and small (S) RNA segments^18^. The L segment encodes the viral RNA-dependent RNA polymerase (RdRp) L protein. Each negative sense RNA segment must first be transcribed into a positive sense messenger RNA (mRNA) by the viral RdRp. The M segment encodes a polyprotein precursor that is proteolytically processed into the Gn and Gc envelope glycoproteins, and the S segment encodes the nucleocapsid protein (N), which binds and packages viral RNA and contributes to viral replication and immune evasion^18-19^. The ANDV S segment also encodes a nonstructural protein, NSs, which antagonizes type I interferon induction by suppressing MAVS-dependent signaling^20^. ANDV and HTNV share (L,∼69%; M,∼65%; S,∼75%) and (RdRp,∼68%; Gp,∼55%; N, ∼65%) nucleic acid and amino acid sequence identity, respectively. Differences in viral protein expression across Old and New World hantaviruses during infection and the extent to which sequence variation enables differential hijacking of endothelial signaling networks has not been reported.

Global multi-omic approaches are well suited to define viral transcript and protein expression and map the host signaling networks differentially engaged by each virus. Transcriptomic studies have shown that hantavirus infection induces antiviral gene programs in endothelial cells, including the production of interferon stimulated genes (ISGs)^21,22^. However, RNA abundance alone cannot capture the full regulatory landscape of infection. Protein abundance defines the functional composition of infected cells, and can conflict with RNA-level measurements, while phosphorylation encodes signaling through kinases, with the ability to rapidly remodel endothelial physiology and structure^23^. Integrated RNA-seq, proteomics, and phosphoproteomics can therefore distinguish transcriptional antiviral programs from post-transcriptional and post-translational remodeling events that directly control endothelial responses to infection.

Here, we generated a time-resolved multi-omic map of primary human endothelial cells infected with ANDV or HTNV by integrating global RNA-seq, global mass spectrometry (MS) proteomics, and MS phosphoproteomics of host and viral transcripts and proteins. We sought to determine whether the two viruses replicate with similar kinetics, express comparable levels of viral proteins, elicit equivalent innate immune responses, and remodel the endothelial cytoskeleton and junctional architecture in the same manner. We additionally sought to identify putative host dependency kinases with antiviral targeting potential. Such data could reveal mechanisms underlying vascular leakage by defining how endothelial architecture and homeostasis are differentially affected by ANDV and HTNV.

## RESULTS

### Hantaviruses exhibit divergent replication kinetics and host responses in primary endothelial cells

Given the tropism of hantaviruses for the vascular endothelium, we focused on characterizing the molecular responses associated with endothelial infection by Old World and New World hantavirus species. Human umbilical vascular endothelial cells (HUVEC) were infected with ANDV or HTNV hantaviruses (MOI = 1) in biological quadruplicate and harvested for bulk RNA sequencing to quantify host and viral transcripts, MS abundance proteomics (AB) to quantify host and viral protein levels, and MS phosphoproteomics (PH) to measure phosphorylation changes on host and viral proteins at 24 and 48 hours post infection (hpi; Fig. 1a). Uninfected mock controls were also included for each time point. Quantification of viral genomes by qPCR revealed similar viral RNA levels at 24 hpi between viruses; however, by 48 hpi, ANDV viral genome levels remained constant, whereas HTNV RNA levels increased by ∼10-fold, consistent with restriction of ANDV and continued replication of HTNV (Fig. 1b). This divergence in replication was independently recapitulated by bulk RNA-seq (Fig. 1c) and proteomics measurements (Fig. 1d), which depicted a more pronounced decline for ANDV transcript and protein levels at 48 hpi.

**Figure 1.**
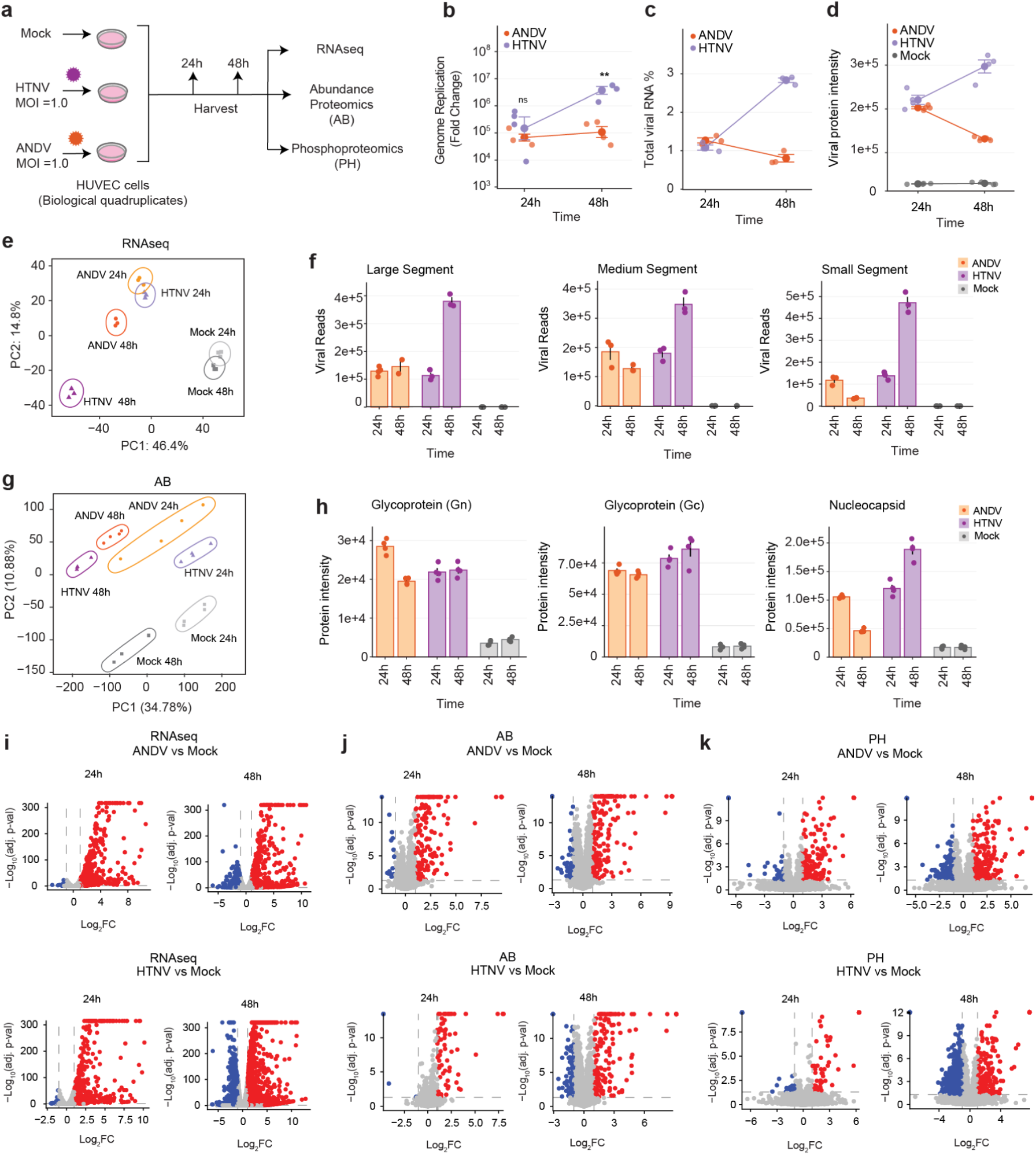
Old and New World hantaviruses induce differing transcriptional and proteomic programs in endothelial cells. (**a**) HUVEC cells were infected with ANDV (MOI 1.0) or HTNV (MOI 1.0) or mock infected (no virus control), and later harvested at 24 or 48 hours post-infection. All conditions were performed in biological quadruplicates. Cell lysates were processed for either RNA-seq to measure changes in gene expression or mass spectrometry (MS) proteomics analysis. For MS analysis, 10% of the total sample was processed for global protein abundance (AB) proteomics analysis and 90% was enriched for phosphorylated peptides to elucidate changes in phosphorylation (PH). diaPASEF method was employed for all MS acquisitions. (**b**) Levels of viral genome replication of ANDV and HTNV in HUVEC cells using strain-specific primer sets. Statistical analysis (Student’s t-test) was performed to compare both viruses: ns, p>0.05; *, p≤0.05; **, p≤0.01; ***, p≤0.001; ****, p≤0.0001. (**c**) Percentage of total viral reads over total gene counts from the bulk mRNA sequencing. (**d**) Sum of all viral protein intensities per virus at different time points post-infection. (**e**) Principal component analysis (PCA) of normalized RNA-seq data from HUVEC cells infected with ANDV (orange), HTNV (purple), or uninfected mock control (gray). (**f**) Number of viral mRNA transcripts for the 3 viral genome segments of hantaviruses. (**g**) Principal component analysis (PCA) of abundance proteomics replicates. (**h**) Quantification of the sum of all viral protein intensities detected from abundance proteomics for each condition over time. (**i**) Volcano plot for RNA-seq, (**j**) abundance proteomics (AB), and (**k**) phosphoproteomics (PH) 24h and 48h after infection. Red dots indicate significantly upregulated (log_2_FC>1 and adj.p-value <0.05) mRNA transcripts, proteins, or phosphorylated peptides for virus-infected cells versus mock-infected cells per time point; blue dots indicate significantly downregulated genes (Log_2_FC< -1 and adj.p-value <0.05).

Within the host, our multi-omics analyses quantified over 23,000 unique RNA transcripts, 77,000 peptides (Extended Data Fig. 1a) mapping to 8700 proteins (Extended Data Fig. 1b), and over 54,000 phosphorylated peptides (∼12,000 with localization probability >0.75; Extended Data Fig. 1d-f) mapping to 4,400 phosphorylated proteins (Extended Data Fig. 1c). Principal component analysis (PCA) of RNA-seq revealed separation of infected versus mock along principal component 1 (PC1) and time point along PC2, with greater separation observed for HTNV at 48hpi (Fig. 1e). PCA for the abundance proteomics and phosphoproteomics analysis also revealed separation by condition (Fig. 1g, Extended Data Fig. 1g).

RNA-seq detected transcripts from all three viral genome segments (Fig. 1f). L, M and S RNA abundance was comparable between ANDV and HTNV at 24 hpi but diverged by 48 hpi (Fig. 1f), which reflected global measurements (Fig. 1b-d). Proteomic analysis detected 3/5 viral proteins—N, Gn and Gc—whereas L remained below background detection and NSs was not detected (Fig. 1h, Extended Data Fig. 1h). N protein abundance closely mirrored S RNA levels, consistent with its expression from the S segment, whereas Gn and Gc abundance remained stable across viruses and time points, suggesting that glycoprotein expression reached a plateau (Fig. 1f,h).

ANDV and HTNV infection dramatically remodeled the transcriptome, proteome, and phosphoproteome of endothelial cells at 24 and 48 hpi. At 24 hpi, HTNV and ANDV induced comparable transcriptional responses, with 572 and 716 genes upregulated and 57 and 103 genes downregulated, respectively. By 48 hpi, HTNV elicited more extensive transcriptomic remodeling than ANDV, with 1999 versus 857 genes upregulated and 1150 versus 467 genes downregulated, respectively (Fig. 1i-k). In contrast, the proteomic responses diverged more strongly at 24 hpi, when ANDV infection induced a dramatic increase in protein abundance, with HTNV increasing the abundance of 80 proteins compared with 163 for ANDV. However, by 48 hpi, HTNV and ANDV elicited comparable responses, with 173 and 181 proteins increased, respectively (Fig. 1j). A similar pattern was observed in the phosphoproteome: ANDV induced more pronounced upregulation of phosphorylation sites at 24 hpi, whereas the HTNV response was comparatively subdued (Fig. 1k). By 48 hpi, the phosphoproteomic responses converged, although HTNV exhibited slightly more extensive decreases in phosphorylation events (Fig. 1k).

### Hantaviruses reprogram immune, cytoskeletal, cell cycle, and mRNA processing pathways in endothelial cells

Gene set overrepresentation analysis (GSOA) of differentially regulated transcripts, proteins, and phosphorylated proteins revealed both viruses elicited a strong innate immune response, which was most evident at the transcript and protein levels, but also at the level of protein phosphorylation (Fig. 2a). In contrast, pathways involved in cytoskeletal organization, cell homeostasis, and RNA metabolism were predominantly dysregulated at the phosphorylation level, despite minimal corresponding changes in transcript or protein abundance, highlighting phosphorylation as a major mechanism of pathway-specific regulation during infection (Fig. 2a).

**Figure 2.**
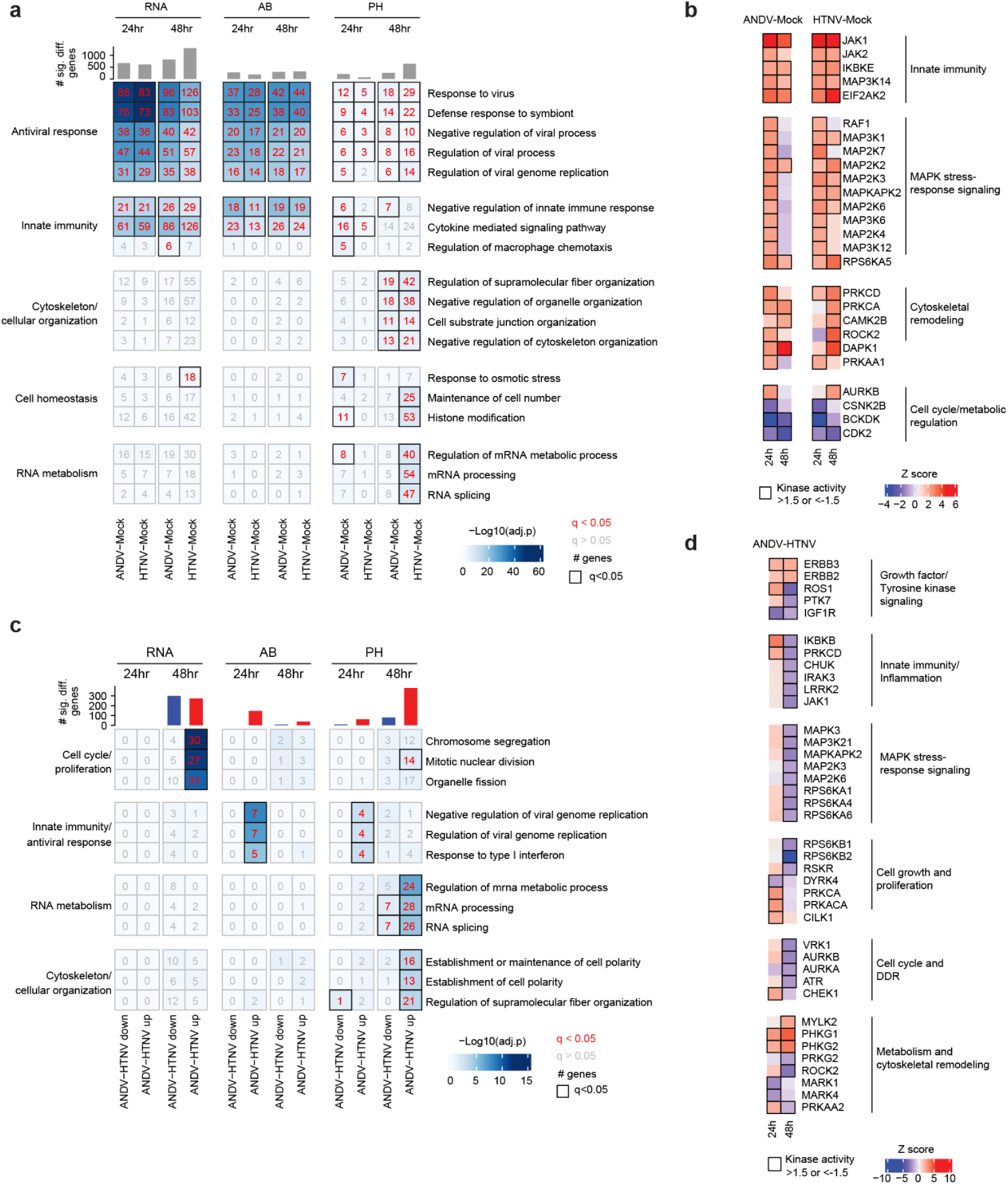
Hantaviruses broadly remodel cellular signaling and host response pathways in endothelial cells. (**a**) Gene Ontology Biological Pathways enrichment analysis of all significantly regulated (abs(Log_2_FC)>1 and adj.p-value <0.05) mRNA transcripts, proteins, and phosphorylated proteins upon infection, separated by time point post-infection and dataset: RNA seq, abundance proteomics (AB), or phosphoproteomics (PH). Numbers in cells indicate the number of genes that match the corresponding term on the right. Red numbers and black squares indicate significantly enriched terms (q<0.05). (**b**) Kinase activities were calculated as described in the Methods section using prior knowledge networks of kinase-substrate interactions. Significantly up- (K>1.5) and downregulated (K<-1.5) kinases are shown inside black squares. (**c**) GSOA results for all significantly up-(Log_2_FC)>1 and adj.p-value <0.05) or down-regulated (abs(Log_2_FC)<-1 and adj.p-value <0.05) mRNA transcripts, proteins, and phosphorylated proteins for the ANDV vs HTNV comparison at 24h and 48h. Top terms were divided into two sets: downregulated (blue) and upregulated (red). (**d**) Kinase activity analysis for the ANDV vs HTNV comparisons at 24h and 48h.

We next inferred temporal changes in host kinase activity from the phosphoproteomics data using prior-knowledge networks of kinase-substrate interactions^24^. This analysis revealed strong activation of innate immune kinases, including JAK1/2, IKBKE, and EIF2AK2 (PKR), consistent with antiviral signaling and, in the case of PKR, stress-induced translational arrest (Fig. 2b). We also observed activation of MAPK signaling at 24 hpi, followed by partial resolution at 48 hpi for ANDV, potentially reflecting reduced viral replication and diminished pathogen-associated molecular pattern (PAMPs) sensing. In parallel, kinases involved in cytoskeletal regulation were activated, whereas cell cycle-associated kinases were broadly inhibited, suggesting impaired cell-cycle progression during infection (Fig. 2b).

Direct comparison of the two viruses revealed enhanced expression of cell cycle transcripts for ANDV at 48 hpi, including pathways involved in chromosome segregation and mitotic nuclear division (Fig. 2c). Relative to HTNV, ANDV infection also depicted higher innate immune protein abundance and phosphorylation at 24 hpi, as well as higher phosphorylation of proteins involved in RNA processing and cytoskeletal organization at 48 hpi (Fig. 2c). This suggests that the two viruses differ in early innate immune antagonist potency and later phosphorylation-dependent remodeling of RNA processing and cytoskeletal networks. Similarly, direct comparison of inferred kinase activities between the two viruses revealed increased activity of IKBKB in ANDV-infected cells at 24 hpi, consistent with enhanced NF-κB signaling, together with activation of ERBB3, ROS1, PRKCD, PRKCA, PRKACA, PRKAA2, CLIK1, CHEK1, and the metabolic regulators PHKG1 and PHKG2 (Fig. 2d). By 48 hpi, ANDV exhibited reduced MAPK stress signaling and diminished innate immune kinase activity, including IKBKB and JAK1 relative to HTNV (Fig. 2d), likely reflecting more effective restriction of ANDV replication and continued HTNV replication at this later stage (Fig. 1b), and higher activity of ERBB2 and ERBB3.

### ANDV elicits a disproportionate innate immune protein response

As noted above, ANDV elicited higher innate immune protein expression at 24 hpi that largely resolved by 48 hpi (Fig. 2c). Although both viruses induced robust expression of ISGs at the transcript and protein levels relative to mock-infected cells (Fig. 3a, Extended Data Fig. 3a,b), direct comparison of the two viruses revealed significantly higher ISG protein expression at 24 hpi in ANDV relative to HTNV (Fig. 3b). Notably, this difference was most evident at the protein level and not at the RNA level (Fig. 3b-e, Extended Data Fig. 3c,d), suggesting that the divergent antiviral responses between the viruses arise primarily through post-transcriptional mechanisms rather than differences in levels of ISG gene activation. However, transcription factor activity analysis, inferred from prior knowledge networks of transcription factor-target interactions and gene level Log2 fold changes, revealed significantly greater activity of IRF1-3, STAT2, RELA/B, and NFKB1 in ANDV compared with HTNV at 24 hpi (Extended Data Fig. 2), depicting a modest signal at the RNA level. These findings further confirm heightened PAMP sensing, interferon signaling, and NFκB activation in ANDV-infected cells.

**Figure 3.**
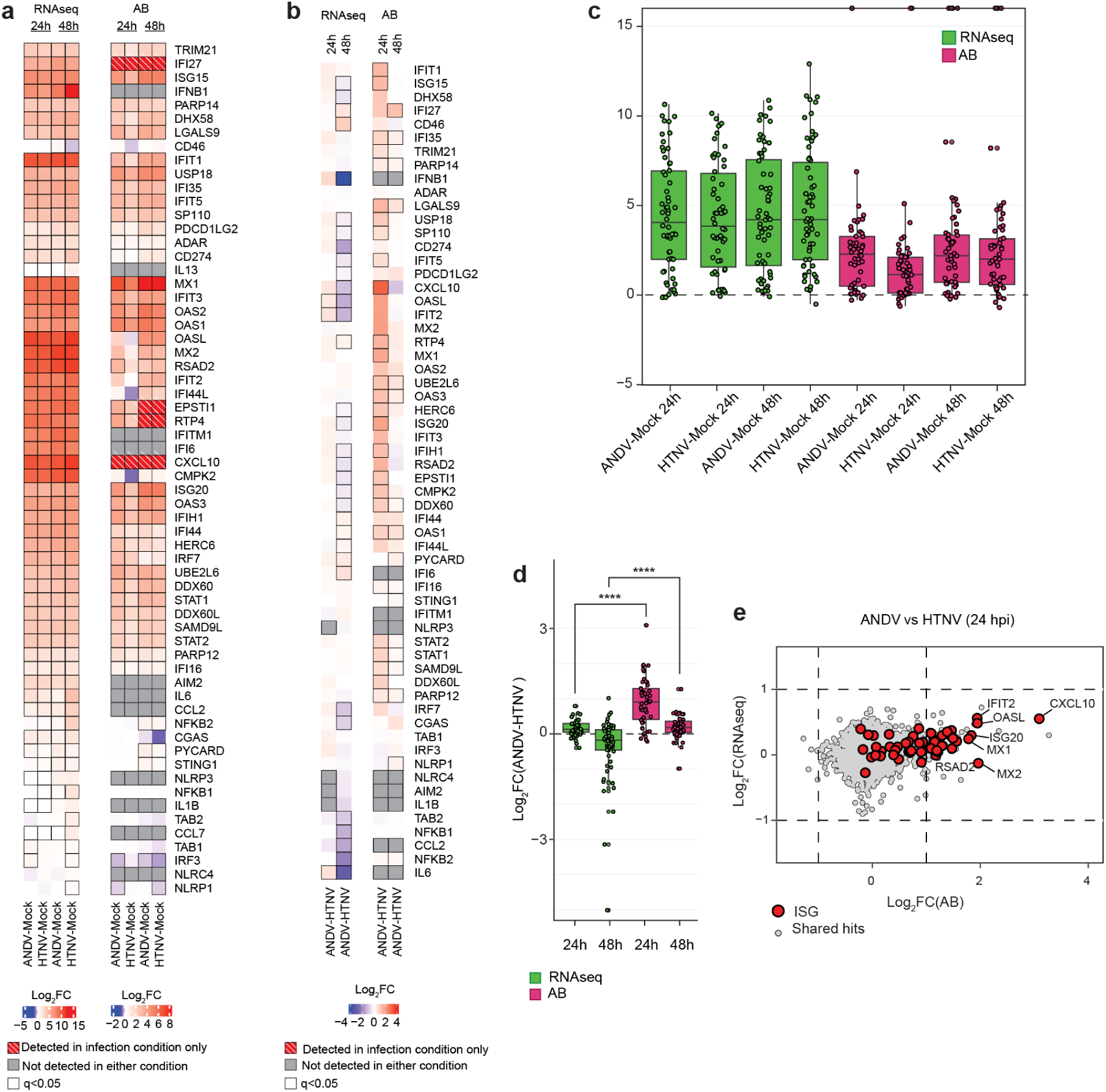
ANDV induces stronger ISG protein, not transcriptional, responses compared to HTNV. (**a**) ISGs mRNA transcript and protein levels for virus-infected HUVEC cells vs mock condition. Black squares indicate significantly regulated terms (q<0.05). (**b**) Comparison of ISGs mRNA transcript and protein levels between both viruses (ANDV vs HTNV). (**c**) Total ISGs log_2_ fold changes in mRNA transcript and protein levels for virus-infected cells vs mock per time point. Statistics determined using Welch’s two sample t-tests. p>0.05; *, p≤0.05; **, p≤0.01; ***, p≤0.001; ****, p≤0.0001. (**d**) ISGs log_2_ fold changes in mRNA transcript and protein levels for ANDV-infected vs HTNV-infected cells at 24 and 48 hpi. Statistics determined using Welch’s two sample t-tests. p>0.05; *, p≤0.05; **, p≤0.01; ***, p≤0.001; ****, p≤0.0001. (**e**) Cross-omics analysis of all shared genes across both the RNA-seq and abundance proteomics datasets at 24 hours post-infection. Red large dots indicate ISGs. Dashed lines indicated boundaries for differentially regulated phosphosites (Log_2_FC <-1 and Log_2_FC > 1). Selected significantly upregulated ISGs on the AB dataset have been labeled.

Enhanced antiviral and inflammatory responses can promote apoptotic cell death, which we sought to evaluate in our system. To determine whether the divergent host responses induced by ANDV and HTNV were associated with differences in apoptosis induction, we examined abundance and phosphorylation changes in apoptosis-related proteins. Both viruses induced phosphorylation changes in these proteins with comparatively little change in protein abundance, although phosphoregulation was more pronounced during ANDV infection (Extended Data Fig. 3e). We next measured caspase-3/7 activity across increasing MOIs. At an MOI of 1, caspase-3/7 activity was comparable between ANDV- and HTNV-infected cells, whereas at an MOI of 10, ANDV induced a modest approximately 3-fold increase (Extended Data Fig. 3f). Thus, despite differential phosphoregulation of apoptosis-associated proteins, the two viruses exhibited relatively limited differences in apoptotic activity under these conditions.

### HTNV infection promotes cytoskeletal dephosphorylation and altered actin organization

Although cytoskeletal pathways were not significantly enriched in the phosphoproteomics dataset at 24 hpi, the two viruses exhibited phosphoregulation of cytoskeletal proteins by 48 hpi (Fig. 2c). Although direct comparison of the viruses appeared to suggest increased phosphorylation of cytoskeletal proteins in ANDV (Fig. 2c), comparison of each virus to mock revealed that HTNV induced a pronounced decrease in cytoskeletal protein phosphorylation at 48 hpi, whereas ANDV exhibited more modest changes relative to mock (Fig. 4a-b). Among cytoskeletal phosphosites decreased at 48 hpi, 21% (36/168) were shared between infections, whereas 79% (132/168) were specific to HTNV (Fig. 4c). Importantly, decreased phosphorylation in HTNV was not associated with concomitant decreases in protein abundance levels (Extended Data Fig. 4a-b), suggesting a signaling-driven response rather than transcriptional protein stability-driven remodeling of endothelial structural networks. Given that HTNV genome replication was elevated relative to ANDV at this later time point, these phosphorylation changes may reflect a direct response to ongoing viral replication.

**Figure 4.**
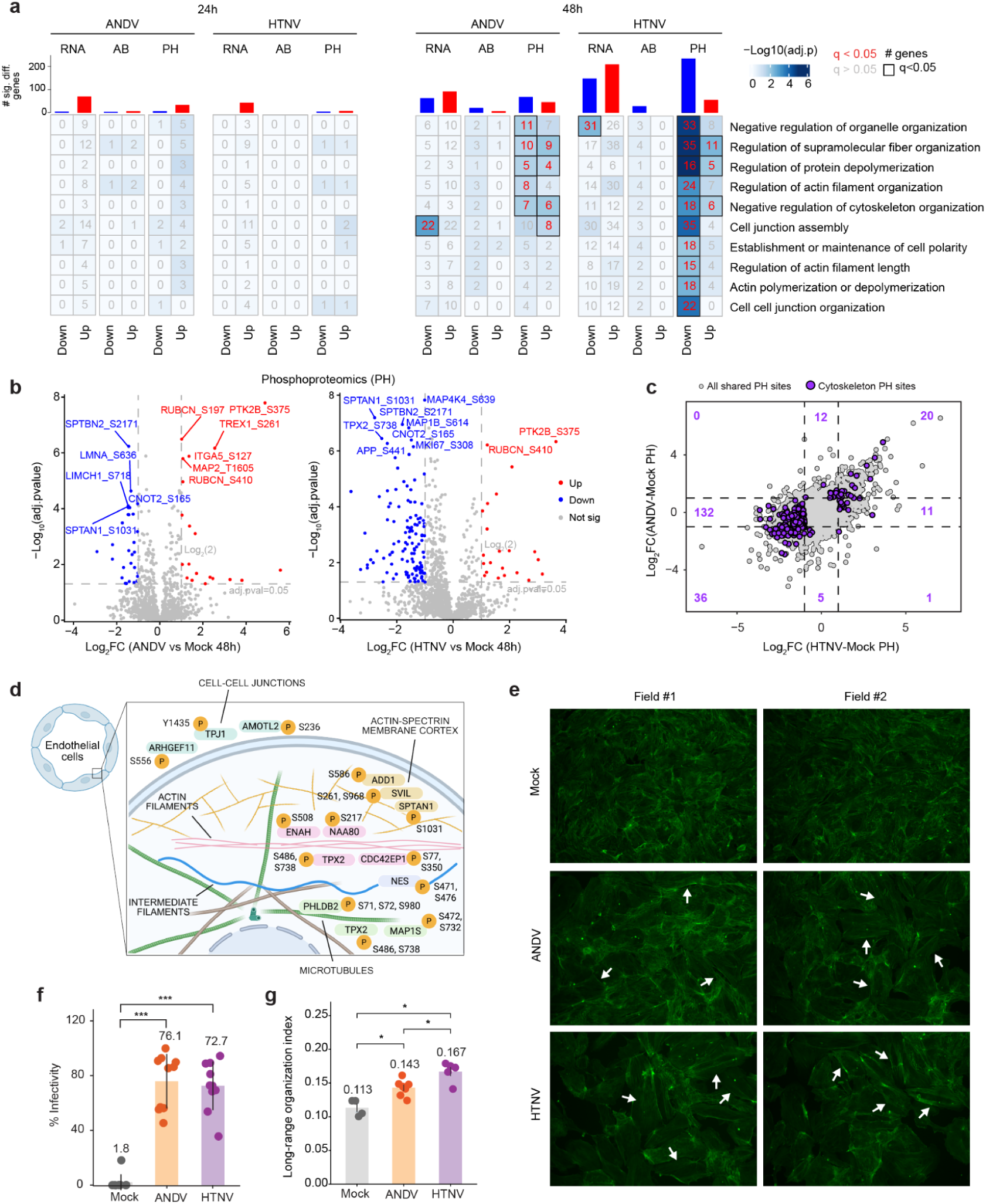
HTNV remodeling of junctional and cytoskeletal proteins is associated with widespread dephosphorylation events. (**a**) GSOA analysis for top terms related to cell morphological changes significantly regulated in the phosphoproteomics data for the ANDV vs HTNV condition at 24h and 48h. Downregulated (blue) and upregulated (red) bar plots quantify the number of significantly changing genes per condition. (**b**) Volcano plots of all unique phosphosites from proteins that match any of the top 5 GOBP terms (**a**) at 48 hpi for virus-infected vs mock-infected comparisons. Significantly upregulated phosphosites (Log_2_FC > 1 & adj.pvalue <0.05) are shown as red dots, some of which are labeled. Significantly downregulated phosphosites (Log_2_FC <-1 & adj.pvalue <0.05) are shown as blue dots. (**c**) Scatterplot of all unique phosphosites that were consistently identified in both ADNV vs Mock and HTNV vs Mock comparisons at 48hpi. Phosphorylation sites on proteins that match the top 5 GOBP terms from (**a**) are highlighted and quantified (purple dots). Dashed lines indicate boundaries for differentially regulated phosphosites (Log_2_FC <-1 and Log_2_FC > 1). (**d**) Schematic overview of phosphorylation events on proteins involved in major types of filaments and cell junctions in endothelial cells upon infection by Hantaviruses. (**e**) Immunofluorescence analysis of F-actin organization in ANDV- or HTNV-infected cells (MOI 10, 48 hpi) in HUVEC cells. Representative images show F-actin filaments (green). White arrows point to elongated F-actin filaments. (**f**) Percentage of positive cells for viral Nucleocapsid in Mock-, ANDV-, or HTNV-infected HUVEC cells at 48 hpi. Quantitative data is presented as the mean ± standard deviation. Statistical analysis was performed using ANOVA, followed by Tukey’s post hoc test. Not significant, ns; *, p≤0.05; **, p≤0.01; ***, p≤0.001; ****, p≤0.0001. (**g**) Long-range F-actin organization index across different conditions (Mock, n=4; ANDV, n= 7; HTNV, n =5). The index measures persistent spatial organization and directionality compatible with actin ridge signal along the local fiber tangent. A higher index value indicates more spatially persistent F-actin organization. Statistical analysis was performed using the Kruskal-Wallis test followed by the Mann-Whitney U test and Holm correction. *, p< 0.05.

HTNV infection was broadly associated with decreased phosphorylation of proteins involved in cell junction assembly, supramolecular fiber organization, actin filament organization, and maintenance of cell polarity (Fig. 4a). Phosphoregulated proteins at 48 hpi mapped to the actin-spectrin membrane cortex, actin filaments, microtubules, intermediate filaments, focal adhesions, and cell-cell junctions, suggesting coordinated remodeling of the structural systems that maintain endothelial shape, adhesion, and barrier integrity (Fig. 4d). Among the phosphosites identified, TAGLN2 S163 and TPX2 S486 have the strongest site-specific functional support. Phosphorylation of TAGLN2 at S163 reduces its association with filamentous actin (F-actin), thereby increasing actin-filament dynamics and cell motility and potentially weakening the cortical actin network that supports endothelial junctions^25^. Phosphorylation of TPX2 at S486 stabilizes the protein and promotes cytoskeletal remodeling and cell migratory behavior^26^. Together with phosphorylation of proteins involved in junction-actin coupling, including TJP1^27^, AMOTL2^28^, and ARHGEF11^29^, and focal-adhesion signaling through PTK2B^30^, these biochemical events could promote junctional disruption and contribute to vascular leakage phenotypes.

To visually assess dysregulation of actin filaments and cellular architecture, we stained HTNV and ANDV cells with DAPI, an antibody against nucleocapsid, and fluorescently conjugated phalloidin at 48 hpi. Imaging revealed pronounced changes in cell distension and F-actin organization (Fig. 4e). Quantification of nucleocapsid-positive cells confirmed infection of over 70% of cells for each virus (Fig. 4f, Extended Data Fig. 4c-d). We then quantified actin organization by identifying F-actin bundles and measuring how consistently each bundle’s signal and orientation persisted along its local trajectory over distances of 5-100 pixels (see Methods). Measurements across all detected actin structures were integrated into a single long-range F-actin organization index for each image, with higher values indicating more elongated and organized actin bundles. This analysis revealed significant cytoskeletal remodeling, most pronounced in HTNV-infected cells, while ANDV-infected cells exhibited an intermediate phenotype (Fig. 4g).

### ERBB-family kinase inhibitors neratinib and afatinib exhibit antiviral activity against ANDV

ANDV poses an imminent threat to public health because of its ability to spread among humans, which can lead to severe pulmonary vascular leak syndrome and death. Further compounding this burden, ANDV is unique among hantaviruses in its documented capacity for person-to-person transmission. We thus sought out to discover a new antiviral therapeutic effective against ANDV. Our phosphoproteomics data indicated increased activity of signaling by ERBB-family receptors (Fig. 2d), leading us to ask whether this pathway represented a functional host dependency for viral replication. ERBB-family receptors regulate cell survival, membrane trafficking, cytoskeletal organization, and junctional remodeling, making them plausible mediators of viral entry, replication, egress, and endothelial remodeling phenotypes observed during infection.

We evaluated the antiviral impact of neratinib and afatinib, both irreversible pan-HER tyrosine kinase inhibitors targeting EGFR/HER1, HER2/ERBB2, and HER4/ERBB4 that are FDA approved for the treatment of HER2-positive metastatic breast cancers^31^ and EGFR-mutant non-small cell lung cancer^32,33^, respectively. As a control, we included favipiravir, a nucleoside analog that inhibits the viral RNA-dependent RNA polymerase. Favipiravir has been approved in some countries for influenza treatment and our group previously found it to be effective against ANDV^34,22^. HUVECs were pre-treated with neratinib (HKI-272; 10 µM), afatinib (BIBW 2992; 10 µM), favipiravir (T-705; 10 µM), or vehicle control for 2 hours following infection with ANDV (MOI=1.0) or mock-infected for 48 hours. At 48 hpi, immunofluorescence staining for the ANDV nucleocapsid protein revealed significantly reduced viral antigen following neratinib, afatinib, and favipiravir treatment compared to vehicle-treated cells (Fig. 5a-b), in the absence of pronounced cytotoxicity (Extended Data Fig. 4e). Furthermore, qPCR targeting the viral genome revealed a significant reduction following treatment with each ERBB inhibitor (Fig. 5c), although the effect was less pronounced than that observed by immunofluorescence (Fig. 5a-b). This discrepancy suggests that viral genome replication may persist to some extent despite reduced viral protein expression, or that residual defective viral genomes remain detectable, which was less the case for favipiravir. The molecular structures of afatinib and neratinib are shown in Fig. 5d-e, highlighting their kinase-binding scaffolds and electrophilic warheads that enable irreversible covalent inhibition of HER-family kinases^35^.

**Figure 5.**
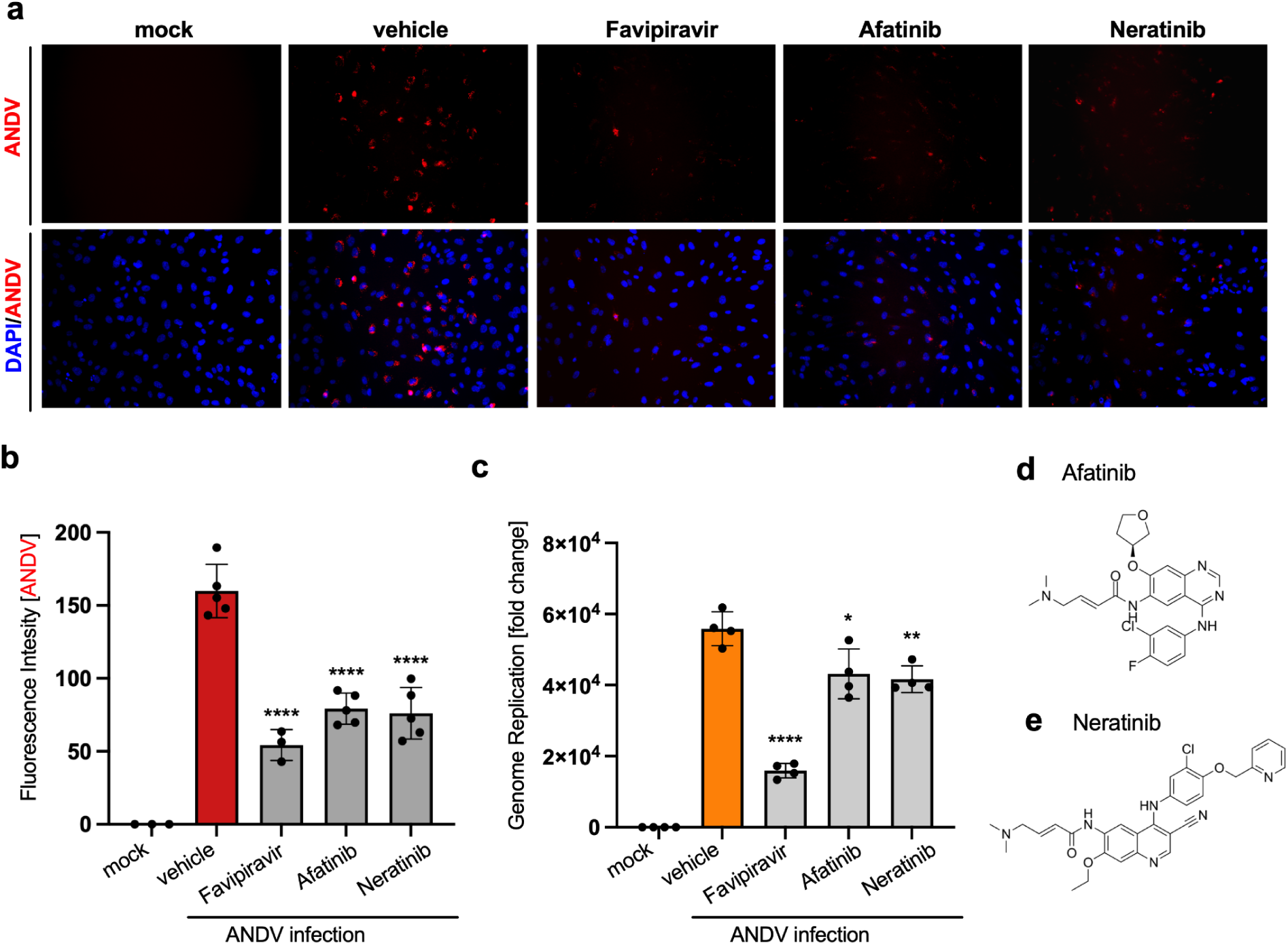
ERBB-family kinase inhibitors neratinib and afatinib exhibit antiviral activity against ANDV. (**a**) Immunofluorescence analysis of HUVEC cells pre-treated with favipiravir (T-705, 10 uM), afatinib (BIBW 2992; 10 µM), neratinib (HKI-272, 10 uM), or vehicle control (DMSO) for 2 hours followed by infection with ANDV at MOI 0.1 for 48 hours post-infection. Cells were tested for the presence of ANDV viral antigen (nucleocapsid protein, N) by standard immunofluorescence techniques. (**b**) Quantification of fluorescence intensity from ADNV nucleocapsid staining channel from immunofluorescence assay in **a**. (**c**) Fold change of viral genome levels by qPCR relative to housekeeping gene. (**d-e**) Chemical structures of afatinib and neratinib.

## DISCUSSION

Our integrated analysis reveals that ANDV and HTNV follow distinct virus-host trajectories in primary endothelial cells, despite comparable infection levels early at 24 hpi. HTNV viral RNA levels continued to rise, consistent with sustained productive replication, whereas ANDV replication stalled at 48 hpi. Despite equivalent viral genome and protein levels, ANDV elicited a stronger early antiviral protein response at 24 hpi, a difference not reflected as prominently at the RNA level, followed by a remodeling of phosphorylation of cytoskeletal and junction proteins at 48 hpi. We additionally found the inhibition of ERBB receptors to be antiviral, suggesting a proviral role for epidermal growth factor (EGF) signaling.

Previous studies have reported distinct tissue tropism for ANDV and HTNV, with ANDV preferentially affecting the lungs and HTNV, the kidneys^22^. However, both viruses converge on the infection of endothelial cells. We find species-specific differences in the timing and nature of the endothelial host response to infection. In endothelial cells, HTNV appears to most strongly antagonize the host innate immune response early, which permits its replication to higher titers, whereas ANDV induces a more pronounced host innate immune response earlier, which leads to the restriction of viral replication at later stages. Enhanced inflammatory signaling can destabilize the cytoskeleton and promote endothelial dysfunction^36–38^. This distinction could explain why ANDV exhibits reduced replication but still produces severe vascular pathology. HTNV, by contrast, may achieve sustained replication through more effective suppression of host innate immune protein translation, with vascular dysfunction emerging through a different balance of viral persistence, inflammation, and tissue-specific endothelial responses^39^.

Although transcription factor activity analysis revealed differences in type I interferon and NFκB programs, no ISG transcripts were significantly different between ANDV and HTNV at 24 hpi. The divergence emerged primarily at the protein level, indicating only a modest transcriptional distinction in their early innate immune responses. The discordance between antiviral RNA and protein responses points to regulation downstream of transcription, including differences in mRNA translation, protein stability, or both. One explanation for this observation is that HTNV more effectively suppresses the translation or accumulation of ISG proteins despite comparable induction of ISG transcripts. This finding is particularly intriguing given previous studies implicating the hantavirus nucleocapsid (N) protein in modulation of host translation by directly associating with host translational machinery^18-19^. HTNV N protein may redirect host translation by functionally substituting for components of the eIF4F initiation complex, thereby favoring viral mRNA translation^18^. These effects are more likely driven by sequence dependent differences in N protein function and virus-host protein-protein interactions than by differences in N abundance, which was comparable between viruses at 24 hpi.

Finally, our identification of ERBB-family signaling as a host dependency factor for ANDV reveals a potentially druggable pathway supporting hantavirus infection. Both neratinib and afatinib significantly reduced viral antigen and genome levels, demonstrating antiviral activity against ANDV. These drugs are irreversible pan-HER kinase inhibitors that target EGFR/HER1, HER2/ERBB2, and HER4/ERBB4 through an electrophilic acrylamide moiety that covalently engages the kinase domain, providing sustained suppression of receptor signaling. ERBB-family receptors regulate membrane trafficking, endocytosis, intracellular signaling, and cell survival, raising several possibilities for how their activity may support the hantavirus life cycle. ERBB signaling could facilitate viral entry or intracellular trafficking, establish a cellular state permissive for viral replication, or promote later stages of virion assembly and egress. Because inhibitors were administered before infection, our experiments do not distinguish among these possibilities, and time-of-addition and genetic perturbation studies will be required to define the specific stage at which ERBB signaling is required. Nevertheless, the antiviral activity of two mechanistically related ERBB inhibitors, together with the increased ERBB-family kinase activity inferred from our phosphoproteomics data, supports a functional role for this signaling axis in ANDV infection and identifies an FDA-approved drug class with potential for therapeutic repurposing.

Together, integration of our multi-omics datasets revealed distinct layers of host regulation that differentiate ANDV and HTNV infection. Combined RNA-seq and proteomic analyses showed that divergent innate immune responses were more pronounced at the protein than transcript level, while phosphoproteomics uncovered widespread cytoskeletal dephosphorylation associated with structural remodeling. Kinase activity inference further identified ERBB-family signaling as a host dependency, leading to the discovery that pharmacologic ERBB inhibition suppresses ANDV infection. Thus, integrating complementary omics approaches not only revealed virus-specific mechanisms of host remodeling but also identified a therapeutically actionable pathway for antiviral drug development.

## METHODS

### Ethics Statement

This study was performed in strict accordance with the recommendations of UCLA. All ANDV, and HTNV live virus experiments were performed at the UCLA BSL-3 High Containment facility.

### Cells

Primary, pooled Human Umbilical Vein Endothelial Cells (HUVECs) were obtained from the American Type Culture Collection (ATCC, PCS-100-022) and cultured in Vascular Cell Basal Media (PCS-100-030, ATCC), supplemented with Endothelial Cell Growth Kit-VEGF (PCS-100-041) and Penicillin-Streptomycin-Amphotericin B Solution (PCS-999-002, ATCC). The cells were maintained in a humidified environment at 37°C, 5% CO_2_, and subcultured following the manufacturer’s recommended specifications. Primary Human Pulmonary Endothelial Cells were used for experiments within passages 2–5.

### Viruses

ANDV (Chile-9717869 strain) was obtained from Dr. Heinz Feldmann from the NIH/NIAID, and HTNV (Fojnica strain) was obtained from BEI Resources. The ANDV Chile-9717869 strain was provided by Dr. Connie Schmaljohn, U.S. Army Medical Research Institute of Infectious Diseases, Ft. Detrick, MD, to Dr. Heinz Feldmann. The Chile-9717869 strain of ANDV was first isolated from an infected *Oligoryzomys longicaudatus* rodent in 1997^40^. The HTNV Fojnica strain was isolated in *Apodemus flavicollis* (yellow-necked wood mouse) in the Fojnica region of former Yugoslavia (now Bosnia and Herzegovina) in 1989^41^. We have amplified these viruses once in Vero E6 cells. Viral stocks were made from cell-free supernatants collected at Day 6 post-infection, aliquoted, and stored at -80°C. We verified the sequences of the S and M genomic segments and confirmed their sequence to respective strains. The virus titer was measured in Vero E6 cells with the established 50% tissue culture infectious dose (TCID50) assay.

### HUVEC cell infection for RNA-seq analysis

HUVECs were plated at 4x10^6^ cells per 10-cm dish or 1x10^5^ cells per well using a 48-well plate. Viral inoculum of ANDV (Chile-9717869 strain), and HTNV (Fojnica strain), was added onto the cells at a multiplicity of infection (MOI) of 1 using serum-free base media. After 1 hour of incubation at 37°C with 5% CO_2_, the inoculum was replaced with endothelial cell-type media. Cells at 10-cm dishes were then lysed using 8 M Urea (Promega) solution at 24 and 48 hours post-infection and stored at -80C. 48-well cells were fixed at selected time points with 4% PFA for 30 minutes at room temperature and subsequently washed three times with ice-cold PBS for IFA staining. ( For RNA samples, the cells were lysed using Trizol for 5 minutes. Vials containing Trizol lysates were stored at -80°C prior to RNA isolation.

### Drug Compounds and Infections

The compounds tested were obtained from Medchem Express. All compounds were provided as lyophilized and were then reconstituted in Nuclease-Free water (Invitrogen) or DMSO. Compounds were then aliquoted and stored at either -80°C or room temperature in dry conditions. For drug studies, indicated drugs were added in HUVECs 2 hours prior to ANDV infection (MOI 0.1). The cells were then fixed with 4% PFA for 30 minutes at 48 hpi for IFA analysis.

### Immunohistochemistry

The fixed cells were washed three times with 1x PBS and permeabilized by incubating in blocking buffer (0.3% Triton X-100, 2% BSA, 5% Goat Serum, 5% Donkey Serum in 1X PBS) for 1 hour at room temperature. For immunostaining, cells were incubated overnight at 4°C with each primary antibody, then washed with 1X PBS three times and incubated with respective secondary antibody [Goat anti-Mouse IgG (H+L) Cross-Adsorbed Secondary Antibody, Alexa Fluor 555 (Thermo Fisher Scientific, CAT#A-21422); Goat anti-Rabbit IgG (H+L) Cross-Adsorbed Secondary Antibody, Alex Fluor 488 (Thermo Fisher Scientific, CAT#A-11008] for 1 hour at room temperature. The cell nuclei were stained with DAPI (4’,6-Diamidino-2-Phenylindole, Dihydrochloride) (Life Technologies) at a dilution of 1:5000 in 1X PBS. Image acquisition was done using Leica DM IRB fluorescent microscopes.

### Image Analysis

Number of cells per field and percentage infectivity were determined using the built-in scripting language Macro (.ijm) on Fiji (ImageJ), which can be found on the Bouhaddou lab github account. The long-range F-actin organization index was determined using the green channel (Phalloidin staining, F-actin) of each TIFF image using a track-free, direction-aware analysis using Codex. Specifically, images were first independently normalized between their 1st and 99.8th intensity percentiles to minimize differences in overall fluorescence and were corrected for uneven background lighting by subtraction of a Gaussian-smoothed image (sigma, 18 pixels). Bright ridge-like actin structures were enhanced using a multiscale Meijering filter evaluated at scales of 1-4 pixels. Local actin orientation and anisotropy were estimated from the structure tensor (Gaussian sigma, 2 pixels); because the tensor’s dominant gradient is perpendicular to a bright ridge, its orientation was rotated by 90° to obtain the actin tangent. Pixels exceeding the image-specific Otsu threshold for ridge response and having structure-tensor coherence of at least 0.25 were retained as ridge seeds. From each seed, ridge strength, local coherence, and axial orientation agreement with the seed orientation were sampled in both directions along the local tangent at 5-pixel intervals from 5 to 100 pixels. Directional agreement was evaluated with a 180°-periodic orientation score, and sampled support was normalized to the ridge strength at the seed. The ridge-weighted mean support-versus-distance curve was integrated from 5 to 100 pixels and divided by the interval length to obtain one long-range tangent-support index per image. Higher values indicate more persistent spatial organization along the actin-fiber direction but do not represent the physical length of an individually traced filament. Images were treated as the statistical units. Differences among conditions were assessed using a Kruskal-Wallis test followed by exact two-sided Mann-Whitney U tests for all three pairwise comparisons, with Holm correction within the endpoint.

### Apoptosis assay

Apoptosis in HUVEC cells infected by ANDV or HTNV at different MOIs (0.5, 1, 4, and 10) was determined using the Caspase-Glo 3/7 assay according to manufacturer’s protocols (Cat.# G8090; Promega, Madison, WI, USA). The caspase activity was measured using Glomax Discover Microplate Reader (Promega). The fold change relative to mock was determined using 4 biological replicates.

### RNA sample preparation and RT-qPCR analysis

RNA was extracted from various virus-infected cells using the RNA Mini Kit (BioRad) according to the manufacturer’s guidelines. Quantification of RNA was performed with a NanoDrop 2000 Spectrophotometer (Thermo Fisher Scientific). For cDNA synthesis, 1 µg of RNA was used alongside random hexamer primers and the SuperScript III reverse transcriptase kit (Thermo Fisher Scientific). qPCR was then carried out using either SYBR Green ROX Supermix (Life Technologies) on an Applied Biosystems QuantStudio 12K Flex RT-PCR system (Thermo Fisher Scientific) or SSOAdvanced Universal SYBR Green Supermix (Bio-Rad) with a CFX384 Touch RT-PCR detection system (Bio-Rad). The reactions were conducted in 10 µL volumes within a 384-well plate, with thermal cycling conditions starting with an initial denaturation at 95°C for 30 seconds, followed by 40 cycles of 95°C for 15 seconds and 60°C for 60 seconds. A melt-curve analysis was performed from 65°C to 95°C, with increments of 0.5°C every 2-5 seconds. The 2-ΔCT method was used for transcript quantification, normalizing against glyceraldehyde-3-phosphate dehydrogenase (GAPDH) CT values^42^. Baseline mRNA levels in mock-treated cells were set to 1, and fold changes in infected cells were calculated relative to this baseline. Details of qPCR primer sequences for the target mRNA transcripts can be found in Table S1.

### RNA sequencing and data analysis

Total RNA samples were prepared as described above. For each experimental condition, three replicate RNA samples were submitted to the UCLA Technology Center for Genomics & Bioinformatics (TCGB) for RNA sequencing analysis. Library preparation, sequencing, and RNA-Seq data analysis were performed as described^43,44^ with minor modifications. In summary, libraries were prepared with the KAPA Stranded mRNA-Seq Kit, followed by second strand synthesis converting the cDNA:RNA hybrid to double-stranded cDNA (dscDNA) and incorporating dUTP into the second cDNA strand. cDNA generation was followed by end repair to generate blunt ends, A-tailing, adaptor ligation, and PCR amplification. Different adaptors were used for multiplexing samples in one lane. Sequencing was performed on Illumina NovaSeq X Plus for a paired-end 2x50 bp run. Data quality checking was done on Illumina SAV. Demultiplexing was performed with Illumina Bcl2fastq v2.19.1.403 software. Partek Flow^16^ was used for all data analysis. Illumina reads from all samples were aligned to the combined human (GRCh38) and viral reference genomes (Andes virus: AF291702.1, AF291703.2, AF291704.5; Hantaan virus: NC_005218.1, NC_005219.1, NC_005222.1) using STAR v2.7.11b ^9,17^, and Ensembl transcripts release GRCh38.107 GTF was used for gene feature annotation. Subsequently, the read counts per gene were quantified.

Differential gene expression analysis was performed using DESeq2 v1.42.1^45^ in R v.4.3.2^46^. The median of ratios method was used to normalize expression counts for each gene across all samples studied. DEGs were considered if they were supported by a false discovery rate (FDR) p < 0.01 or pad 0.01 & FC 1. Unsupervised principal component analysis (PCA) was performed using DESeq2 in R v.4.3.2^46^. We deposited RNA-seq data to the NCBI GEO under the accession number GSE232641.

### Proteomics sample processing

Samples were lysed in 8 M Urea (Promega) in LoBind tubes (Eppendorf) and stored on ice until sonication. Lysed samples were sonicated using a probe sonicator 5x for 10 seconds On and 10 seconds Off at 10% amplitude, and protein was quantified using a bicinchoninic acid assay (BCA). Approximately 200 µg of protein for each sample was used for further processing, starting with reduction and alkylation using a 1:10 sample volume of tris-(2-carboxyethyl) (TCEP) (10 mM final) and 2-chloroacetamide (40 mM final) for 30 minutes each at 23°C with shaking at 1100 rpm. Before protein digestion, the 8 M Urea was diluted 7-fold with 100 mM Tris-HCl (pH 8) to permit the activity of the proteolytic enzyme trypsin. Trypsin (Promega) and Lys-C (Wako) was added at a 1:100 (wt/wt) enzyme-substrate ratio and placed in a thermomixer at 23°C overnight (16 h) with shaking at 1000 rpm. Following digestion, 10% trifluoroacetic acid (TFA) was added to each sample to a final pH of approximately 2-3. Samples were then desalted using a vacuum manifold with Oro-Flex I Polypropylene filter plate w/10 m PE frit 96 well plate (Orochem) with 7 mg HLB Resin (Oasis) for 100 ug of peptide. Each well with resin was activated with 100 uL of 100% acetonitrile (ACN), then equilibrated with 3 x 100 uL of 0.1% TFA. After sample loading, the wells were washed with 3 x 100 uL of 0.1% TFA, and samples were eluted with 2 x 60 uL 50%ACN/0.25% formic acid (FA). Approximately 10% of the desalted peptides were separated for global abundance proteomics analysis and the remaining 90% were used for phosphopeptide enrichment. Both the fractions were then dried by vacuum centrifugation in a SpeedVac (Labconco). For phosphopeptide enrichment, Ti-IMAC beads (Resyn Biosciences) were aliquoted in a 1:2 (w/w) peptide:beads ratio and equilibrated three times with binding buffer (0.1 M glycolic acid in 80% acetonitrile (ACN), 5% TFA). Dried peptide samples were resuspended in 200 μL of binding buffer, added to the equilibrated beads, and incubated for 30 min at 23 °C, 1200 rpm. The unbound fraction was discarded, and beads were washed sequentially once each with 200 μL of the binding buffer, wash buffer 1 (60% ACN, 1% TFA, 200 mM NaCl), wash buffer 2 (60% ACN, 1% TFA), and finally with LC-MS grade water. Enriched phosphopeptides were eluted twice by incubating the beads with 150 μL of 1% (v/v) ammonium hydroxide (Sigma) in LC-MS grade water (Fisher Scientific) for 10 min at 23 °C, 1200 rpm. The eluted peptides were transferred to a new protein LoBind tube containing 50 μL of 10% (v/v) formic acid in LC-MS grade water. Both eluates were pooled and dried followed by plate-based desalting as explained previously and were further resuspended in 0.1% formic acid. A volume corresponding to approximately 500 ng of peptides was injected into the timsTOF HT mass spectrometer (Bruker Daltonics).

### Mass spectrometry proteomics acquisition

Dried peptides were resuspended in 0.1% (v/v) FA in MS grade water (Fisher) and analyzed on a timsTOF HT mass spectrometer, paired with a Vanquish Neo UHPLC system. Mobile phase A consisted of 0.1% (v/v) FA in MS grade water (Fisher), and mobile phase B consisted of 0.1% (v/v) FA in 100% MS grade Acetonitrile (Fisher). The LC was operated in trap-and-elute mode, where the peptides were first trapped onto a PepMap Neo Trap column (5 mm, 100 Å pore size, 5 µm particle size) and then reversed-phase separated using gradients mentioned below on an Aurora Elite C18 reverse phase column (15 cm, 100 Å pore size, 1.5 µm particle size for captive spray, IonOptiks), kept at 50°C using a column oven for Bruker Captive Spray source (Sonation Lab Solutions), and ionized in a CaptiveSpray source (Bruker Daltonics) at 1700 V. For global abundance proteome analysis, the %B gradient used was: 5% to 35% over 37 min at 0.3 µL/min, then to 45% in the next 4 mins, and 60% in the next 1 min, followed by an increase to 95% B over 3 min. For phosphoproteome analysis, the %B gradient used was 3% to 16% over 26 min, 30% in the next 11.5 min, 45% in the next 4 min, and 60% B for 1 followed by an increase to 95% B over 3 min min at 0.3 µL/min flow rate.

For the global abundance data analysis, equal-size windows of 25 Da were designed with an overlap of 1 Da to maximize the precursors ion coverage for further MS/MS. Ions were accumulated and separated in the TIMS analyzer over a mobility range of 0.60–1.60 V·s/cm² using 100 ms accumulation and ramp times. DIA acquisition comprised one MS1 scan followed by eleven dia-PASEF scans covering a mass range of 384.4–1305.4 m/z and an ion mobility range of 0.77–1.41 1/K0. Collision energies were linearly ramped according to ion mobility from 20 eV at 1/K0 = 0.71 to 59 eV at 1/K0 = 1.41. The resulting cycle time was approximately 1.27 s. For phosphoproteome analysis, the raw data was acquired in dia-PASEF mode with variable isolation window widths in the m/z vs ion mobility plane. For dia-PASEF acquisition, ions were accumulated and separated in the TIMS device over a mobility range of 0.60–1.60 Vs/cm². MS/MS spectra were acquired across a mobility range of 0.67–1.33 1/K0 and a mass range of 307.2–1292.2 m/z. Collision energies were linearly ramped as a function of ion mobility from 20 eV at 1/K0 = 0.71 to 59 eV at 1/K0 = 1.41. The estimated mean cycle time was 1.27 s. For the global abundance data analysis, equal-size windows of 25 Da were designed with an overlap of 1 Da to maximize the precursors ion coverage for further MS/MS.

### Mass spectrometry proteomics data search and quantitative analysis

The raw files were processed with Spectronaut (Biognosys, [Update 20.2.250922.92449]) with the directDIA+ (Deep) search algorithm, its in-silico derived DIA analysis. Carbamidomethylation (cysteine) was set as a fixed modification for database search. Acetylation (protein N-term), oxidation (methionine), and phosphorylation (serine, threonine, tyrosine; only for phosphoproteomics dataset) were set as variable modifications. Reviewed human protein sequences (downloaded from UniProt, October 6, 2023) were used for spectral matching. The false discovery rates for the PSM, peptide, and protein groups were set to 0.01, and the minimum localization threshold for PTM was set to zero. For MS2-level area-based quantification, the cross-run normalization option was unchecked (normalization was performed later using MSstats), and the probability cutoff was set to zero for the PTM localization. Quantitative analysis was performed in the R statistical programming language (v.4.4.1)^46^. Initial quality control analyses, including inter-run clustering, correlations, principal component analysis (PCA), peptide and protein counts, and intensities were completed in R. Statistical analysis of phosphorylation and protein abundance changes between exposed and control samples were computed using the R package MSstats (4.12.1)^47^. For protein abundance, all peptides mapping to the same proteins were summarized together using the Tukey’s Median Polish approach. For phosphoproteomics, all peptides containing the same set of phosphorylated sites were summarized together into phosphorylation site groups using Tukey’s Median Polish. For both phosphopeptide and protein abundance MSstats pipelines, MSstats performs normalization by median equalization, imputation was turned Off (set to FALSE), and statistical tests of differences in intensity between conditions were calculated using default settings in MSstats. Specifically, MSstats calculates log2 fold changes as the ratio of averaged (across replicates) protein or phosphopeptide intensities between conditions, uses a Student’s t-test for p-value calculation and the Benjamini-Hochberg method of FDR estimation to adjust p-values.

### GO enrichment analysis

Significantly up and down regulated genes (adjusted p-value <0.05 & abs (Log_2_FC)>1) on the RNA-seq, abundance proteomics, and phosphoproteomics datasets (Supplementary Table 1) were tested for enrichment of gene ontology (GO Biological Process) terms. Gene set over-representation analysis (GSOA) was performed using the enricher function of clusterProfiler package (version 4.21.0)^48^ with default parameters. Gene ontology terms were retrieved from the c5 category of Molecular Signature Database (Human MSigDBv2026.1.Hs) [26]. Top 5 terms (ranked based lowest adjusted p-value<0.05) per condition (virus-infected cells vs Mock-infected cells) per time point per experiment type were selected for the heatmap (Fig. 2a, Supplementary Table 2). Similarly, a comparative GOBP analysis was performed for ANDV-infected cells vs HTNV-infected cells following the same steps described above. GSOA was performed on significantly up- and down-regulated genes separately. For each time point, the top 12 enriched GO terms were selected and subsequently consolidated into broader functional categories to reduce redundancy. The 3 most enriched representative terms within each broader category, as determined by enrichment in the PH dataset, were retained for heatmap visualization (Fig. 2c, Supplementary Table 2). Finally, the enrichment analysis for ANDV-infected cells vs HTNV-infected was further refined to select non-redundant top terms related to cell morphology and structural organization changes (Fig. 4a). The ggplot2 v4.0.2 in R was used to generate figures. The heatmaps were generated using heatmap v2.20.02 in R.

### Kinase activity analysis

Kinase activities were estimated using known kinase-substrate relationships as performed previously^49^. Log_2_ fold changes (Log_2_FC) were calculated per phosphorylation site intensities between conditions, and these values were used to infer kinase activity states using prior knowledge networks of kinase-substrate interactions derived from the Omnipath database^50^. A Z-test from the comparison of fold changes in phosphosite intensity measurements of the known substrates against the overall distribution of fold changes across the sample was performed to infer kinase activities as a -log10(p-value) (Fig. 2b,d). This statistical approach has been previously shown to perform well at estimating kinase activities^24,50,51^.

### Transcription factor activity analysis

Transcription factor (TF) activity was inferred from RNA-seq differential expression data using a footprint-based approach implemented in decoupleR^52^. First, gene-level statistics were calculated for each comparison as a signed significance score. This statistic equals the sign of the Log_2_FC multiplied by the -log10 of the adjusted p-value. Genes with missing or non-finite values were removed. Second, a ranked gene vector was generated for each comparison and used as the input for TF activity inference. We obtained TF regulons from the DoRothEA v2 human transcription factor network^53^, and we filtered the interactions to only include high-confidence edges (confidence levels A-C). Finally, TF activities were estimated using the univariate linear model (ULM) implemented in decoupleR. This method tests whether the known gene targets for each TF are consistently up- or down-regulated in the RNA-seq data. TF scores were independently calculated for each comparison. The resulting TF activity p-values were corrected for multiple hypothesis testing using the Benjamini-Hochberg false discovery rate (FDR) procedure within each comparison.

## DATA AVAILABILITY

In addition, supplementary information is available for this paper. RNA sequencing data shown in this study have been deposited into the NCBI’s Gene Expression Omnibus (GEO Accession Number:GSE342604 and BioProject accession: PRJNA1497032). For reviewers, the token is “axchoqsqhjkrpax”. Abundance proteomics, phosphoproteomics proteomics, and large computational datasets used here were uploaded to the PRoteomics IDEntifications Database (PRIDE) at ebi.ac.uk/pride under project accession “PXD080349”. For reviewers, the token is “ySxoBHB8YjbP”. Confocal images and custom scripts used in this study are available at the Bouhaddou Lab GitHub account: https://github.com/BouhaddouLab.

## ACKNOWLEDGEMENTS

We acknowledge the UCLA AIDS Institute, the James B. Pendleton Charitable Trust, and the McCarthy Family Foundation for their generous support of our research. We would like to acknowledge funding from the National Institutes of Health (NIH-NIGMS R35GM160071 to MB). We sincerely thank Dr. Sylvia Neumann for her assistance with microscopy and helpful discussions. Our trainees were generously supported by the Howard Hughes Medical Institute (HHMI) Gilliam Fellowship (#GT17454 to SKM), Molecular Biology Institute Whitcome Fellowship to SKM, the UCLA-Caltech Medical Scientist Training Program (NIH-NIGMS T32GM008042 to SFB), the David Geffen Medical Student Scholarship to SFB, the Cellular and Molecular Biology T32 at UCLA (NIH-NIGMS T32GM145388 to DMW), the Microbial Pathogenesis Training Grant (MPTG NIH-NIAID T32AI007323 to YD), the Graduate Research Fellowship Program (GRFP) from the National Science Foundation (DGE-2444110 to YD). This study was partially funded by NIH grants R01EY032149 and R01EY036572 to VA.

## EXTENDED DATA AND FIGURES

**Extended Data Fig. 1.**
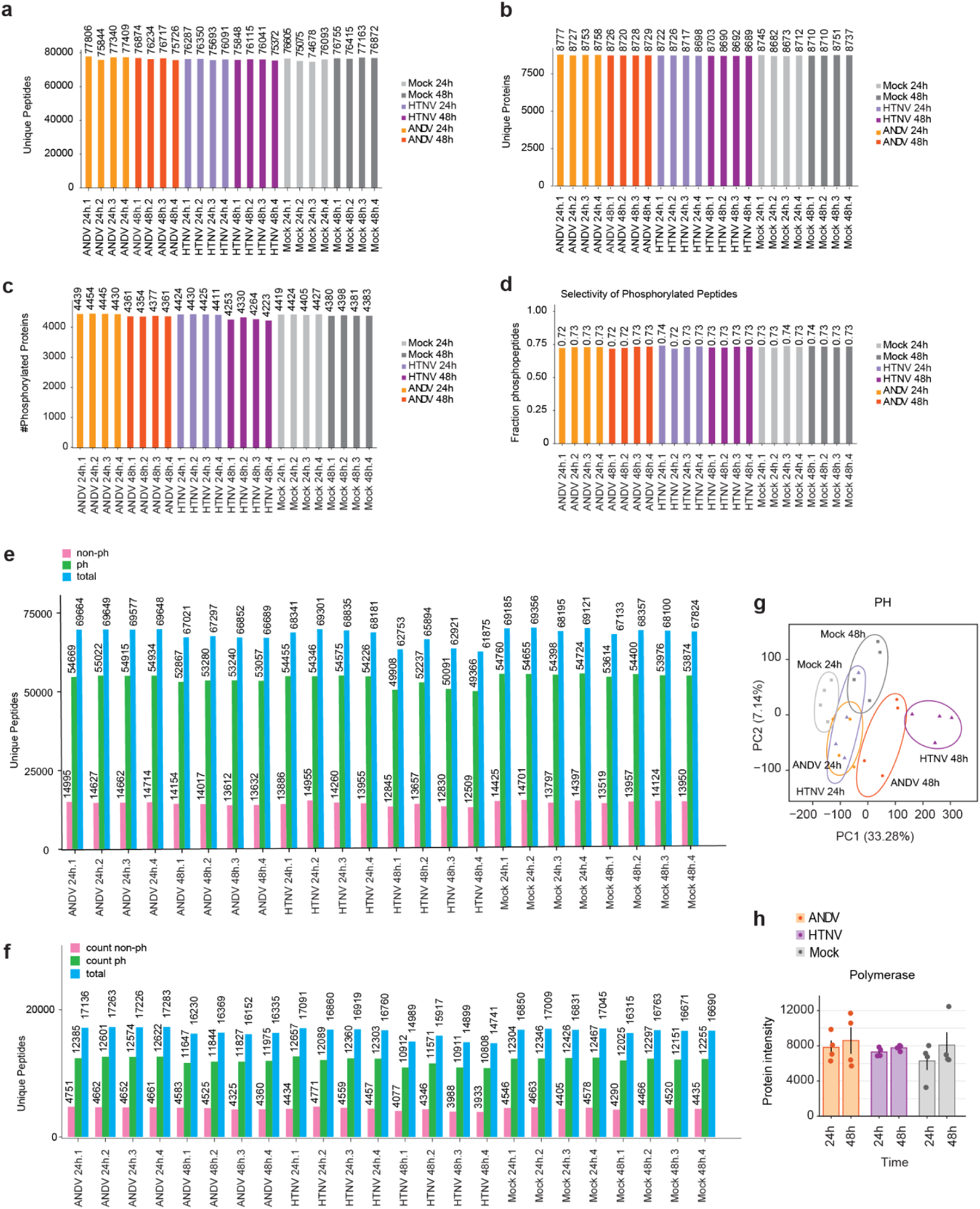
Global omics quality control results. (**a**) Number of unique peptides detected by MS abundance proteomics across different conditions over time. (**b**) Number of unique proteins detected through abundance proteomics across different conditions at indicated time points. (**c**) Number of phosphorylated proteins detected by MS per condition per time point. (**d**) Ratio of detected phosphorylated peptides over total number of detected peptides across conditions and time points. (**e**) Number of unique non-phosphorylated peptides (count non-ph), phosphorylation peptides (count ph), and total peptides (total) detected in the phosphoproteomics analysis. (**f**) Number of high confidence (localization probability >0.75) phosphorylation sites. (**g**) Principal component analysis (PCA) of RNA-seq results highlighting reproducibility among biological replicates and transcriptional differences across treatments (**h**) Quantification of protein intensity for the viral Polymerase protein for different conditions over time.

**Extended Data Fig. 2.**
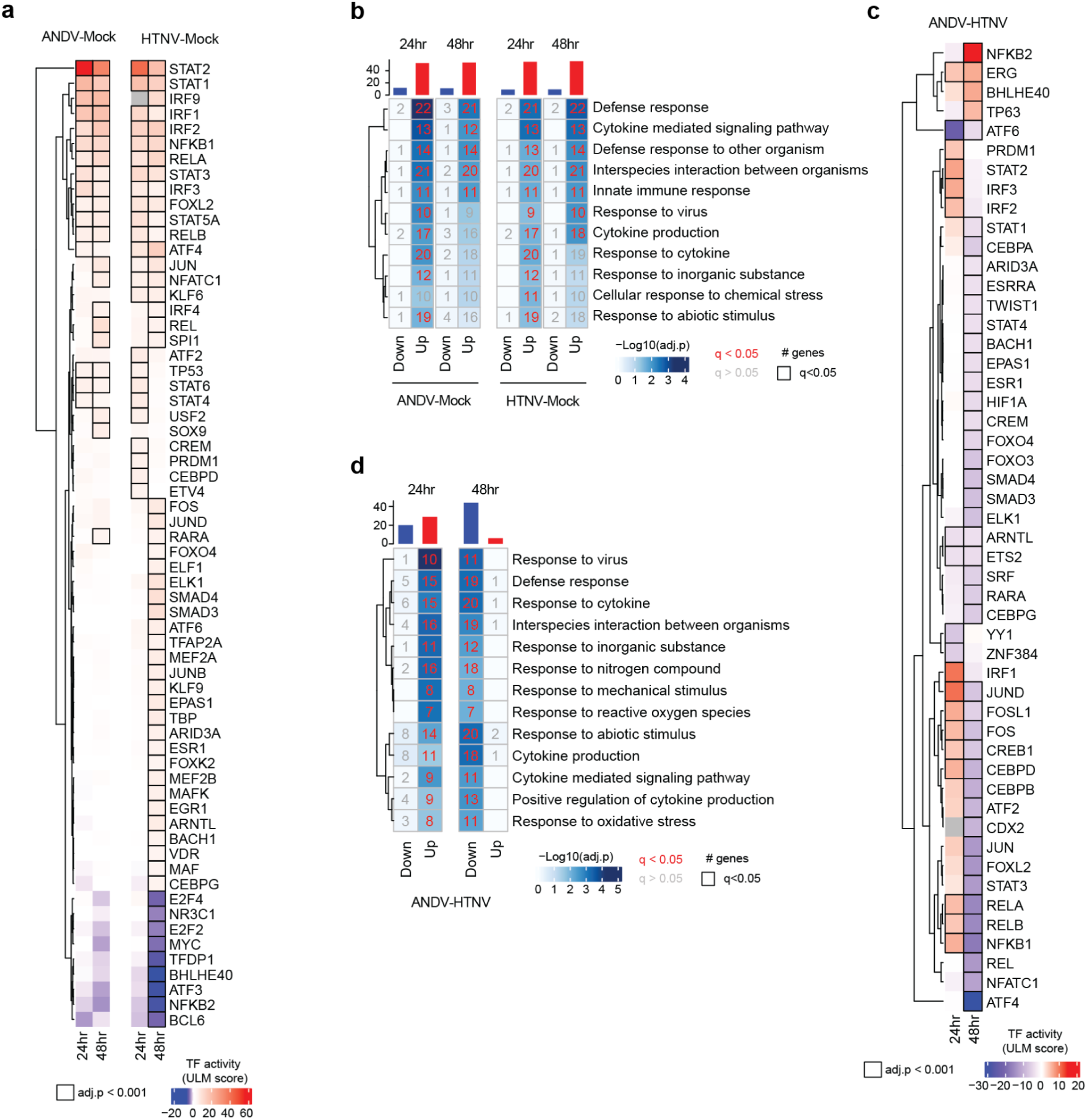
Innate immunity-related transcriptional programs are upregulated in ANDV- and HTNV-infected cells. (**a**) Transcription factor activity inference with univariate linear model (ULM) inferred from RNA-seq differential expression data (Virus-infected cells vs uninfected mock conditions) at different hpi. ULM score is the obtained t-value from the fitted model that predicts observed gene expression based on TF’s TF-gene interaction weights. If ULM score>0, TF is active while if ULM score < 0, TF is inactive. The resulting TF activity p-values were corrected for multiple hypothesis testing using the Benjamini-Hochberg false discovery rate (FDR) procedure within each comparison (adj.p). (**b**) GSOA analysis for significantly regulated TF in the TF activity analysis for the Virus-infected cells vs uninfected mock conditions at 24h and 48h. Top terms were divided into two sets: downregulated (blue) and upregulated (red) bar plots quantifying the number of significantly changing genes per condition. (**c**) Inference of transcription factor activity with ULM using the RNA-seq differential expression data for ANDV vs HTNV comparison at 24 and 48 hpi. (**d**) GSOA analysis for significantly regulated TF for the ANDV vs HTNV comparison at different time points.

**Extended Data Fig. 3.**
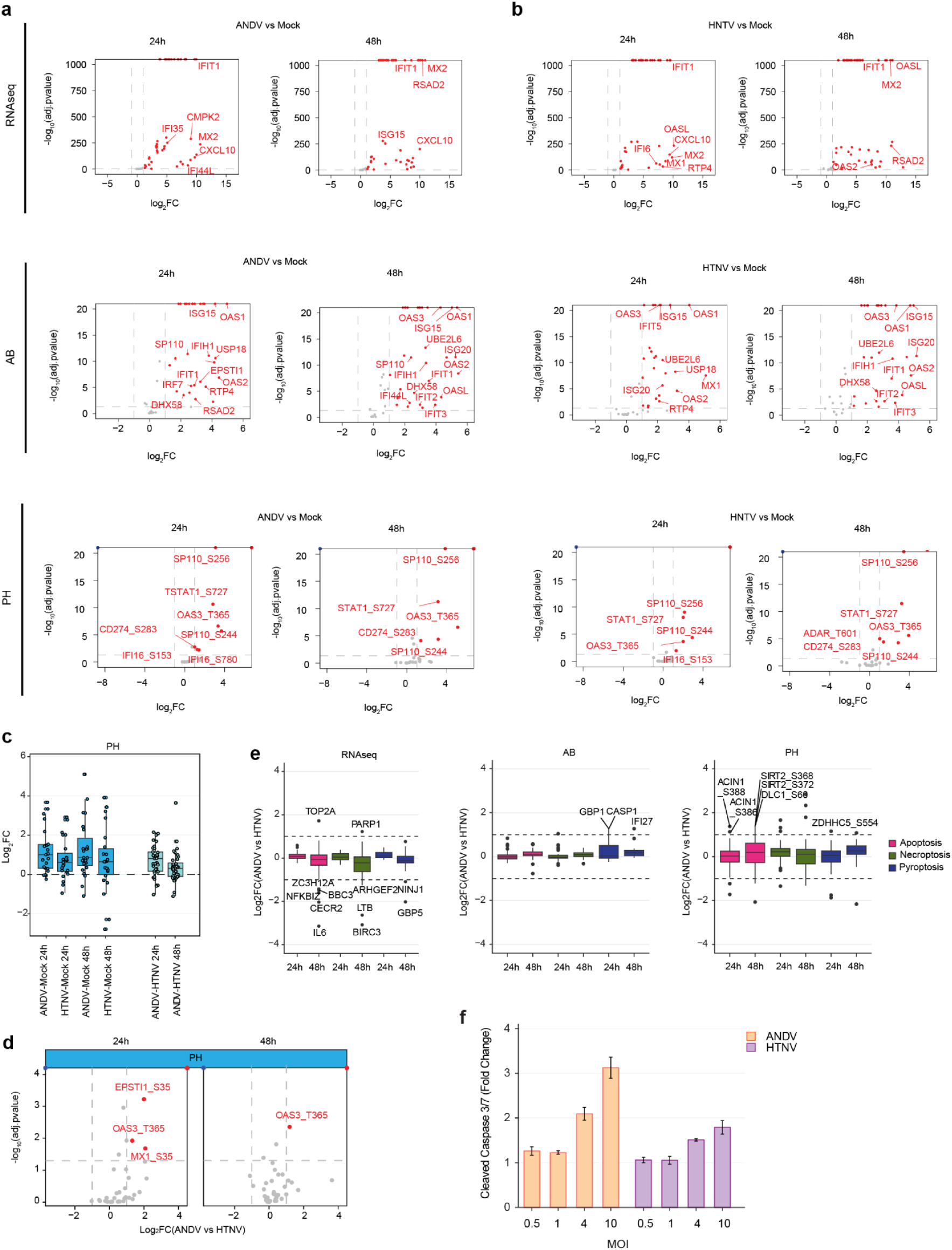
Interferon stimulated gene profiles across RNA-seq, AB, and PH datasets. (**a**) Volcano plots of all ISGs detected in either RNA-seq data (top), AB proteomics (middle), or phosphoproteomics PH (bottom) over time for the ANDV vs Mock comparison. A few significantly upregulated ISGs (red dots) are labeled on the plot. (**b**) Volcano plots of all ISGs detected in either RNA-seq data (top), AB proteomics (middle), or phosphoproteomics PH (bottom) at 24hpi and 48hpi for the HTNV vs Mock comparison. Selected significantly upregulated ISGs (red dots) are labeled on the plot. (**c**) Overall ISG phosphorylation site intensity distribution for virus vs mock and virus vs virus comparisons. (**d**) Volcano plots of all detected phosphorylation sites on ISGs in the PH results. Significantly upregulated (red) phosphorylation sites on ISGs were labeled. (**e**) Temporal changes in genes/proteins/phosphorylation sites involved in programmed cell death pathways in RNA-seq (left), abundance (middle), and phosphoproteomics (right) analyses. (**f**) Fold change of cleaved caspase 3/7 activity in ANDV- or HTNV-infected HUVEC cells versus Mock-infected at different MOIs 48 hpi.

**Extended Data Fig. 4.**
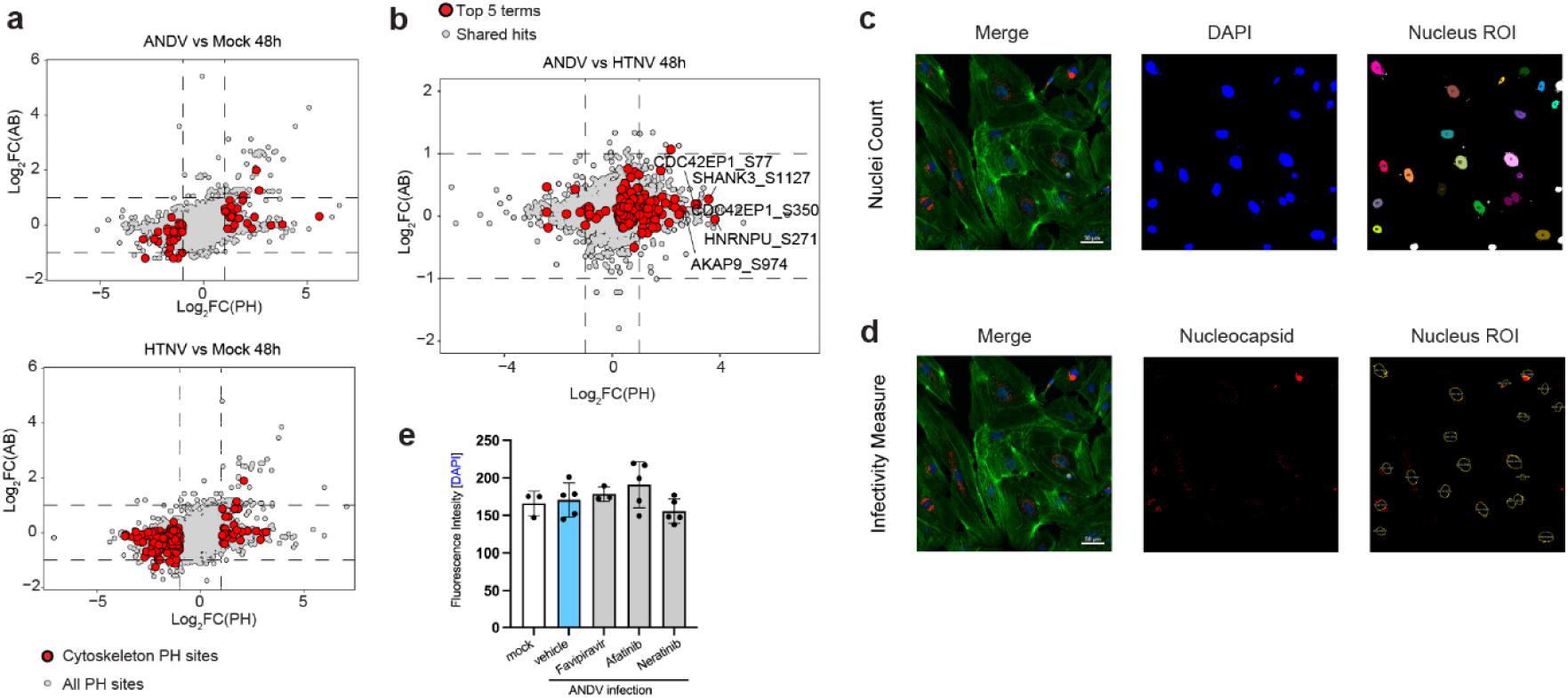
Cellular morphology changes induced by hantavirus infection. (**a**) Scatterplot of all unique phosphosites identified in proteins detected in the abundance proteomics dataset in ANDV vs Mock infected cells (upper panel) or HTNV vs Mock infected cells (lower panel) at 48 hpi. Differentially regulated phosphorylation sites on proteins that match the cytoskeleton remodeling GOBP terms are highlighted (red dots). Dashed lines indicate boundaries for differentially regulated phosphosites (Log_2_FC <-1 and Log_2_FC > 1). (**b**) Comparison between all unique phosphosites identified in both the ANDV- and HTNV-infected samples at 48 hpi. All cytoskeleton-related phosphosites are shown as red dots. Dashed lines indicate boundaries for differentially regulated phosphosites (Log_2_FC <-1 and Log_2_FC > 1). Selected significantly upregulated phosphosites in ANDV-infected cells compared to HTNV are labeled. (**c**) Representative confocal images for the nuclei counting pipeline utilized to quantify the number of nuclei per field of view defining region of interests (ROI) around objects delimited by the DAPI channel. HTNV_003 is shown in this representative panel. Fiji script is available on the Bouhaddou lab github account. (**d**) Representative confocal images for the percentage infectivity approach utilized to calculate percentage of infected cells. ROIs around objects delimited by the DAPI channel (cell nucleus) were first determined. The red channel, which reports on the presence of viral antigen (nucleocapsid protein, N) was overlapped with the ROIs, and the number of cells positive for the red channel were recorded. HTNV_003 is shown in this representative panel. Fiji script is available on the Bouhaddou lab github account. (**e**) Immunofluorescence analysis of cell nuclei (DAPI staining) for HUVEC cells mock-infected or ANDV-infected either treated with vehicle control or ERBB kinase inhibitors

## SUPPLEMENTARY TABLES

**Table S1. qPCR, RNA-seq, Abundance, and Phosphorylation Data, Related to Figure 1**

Contains PCR virus-specific primer sequences and qPCR raw results (PCRData tab). Also, contains all RNA-seq and proteomic datasets of HUVEC cells upon ANDV, HTNV, or Mock infection. Find the full list of mRNA transcripts changes (RNAseqData tab), full list of protein abundance measurements (AbundanceData tab), and phosphorylation sites occurring upon virus infection (PhosphoData tab). Column descriptions are indicated in the first tab.

**Table S2. Enrichments, Related to Figure 2, 3, and 4**

Gene ontology enrichment analyses for significantly changing gene/proteins in all 3 datasets (RNA-seq, abundance proteomics (AB), and phosphoproteomics (PH)) upon ANDV or HTNV infection compared to Mock (Virus vs Mock GSOA tab) or each other (ANDV vs HTNV GSOA tab). Column descriptions are indicated in the first tab.

**Table S3. Kinase Activity Analysis, Related to Figure 3**

Full results of kinase activities for each time point for virus vs mock and virus vs virus comparisons (Kinase Activity Analysis tab). Kinase activities were inferred as a -log10(p-value) of Z-test from the comparison of fold changes in phosphorylation site measurements of the known substrates against the overall distribution of fold changes across the sample. Column descriptions are indicated in the first tab.

**Table S4. Transcription Factor Analysis and Enrichment, Related to Extended Data Figure 3**

Inference of transcription factor activity from RNA-seq differential expression data for all comparisons at different hpi (TF activity tab). Gene ontology enrichment analyses for TFs with statistically significant activity scores (adj.pvalue <0.001) for ANDV- or HTNV-infected samples compared to Mock (GOBP TF activity VvsM tab) or virus-infected samples only (GOBP TF activity VvsV tab).

**Table S5. Cleaved Caspase 3/7 Activity, Related to Extended Data Figure 3**

Raw results and analysis of caspase activity for apoptosis detection in HUVEC cells infected by ANDV or HTNV at different MOIs.

**Table S6. Immunofluorescence Cytoskeleton Remodeling Analysis, Related to Figure 4**

Percentage infectivity calculations for all ten fields of view from confocal images taken for mock, ANDV-, or HTNV-infected (MOI = 1) HUVEC cells 48 hpi.

